# Control of mitotic spindle disassembly through SUMO-targeted ubiquitylation of yeast kinesin-5

**DOI:** 10.1101/2025.11.10.687237

**Authors:** Florence Gaven, Davide Panigada, Didier Portran, Dimitris Liakopoulos

## Abstract

Faithful cell division requires precise control of spindle assembly and disassembly. In budding yeast, spindle breakdown at mitotic exit depends on Cdk1 inactivation and ubiquitin-mediated degradation of spindle stabilizing factors, including the kinesin-5 Cin8 and the crosslinker Ase1/PRC1. Timely degradation of Cin8 by APC^Cdh1^ prevents spindle over-elongation and deformation. We have identified SUMOylation and the SUMO-targeted ubiquitin ligase Slx5 as regulators of spindle disassembly. Cin8 is SUMOylated and interacts with Slx5, suggesting that it is a substrate of Slx5. Deletion of the *SLX5* gene or expression of a SUMOylation-defective Cin8 mutant cause spindle disassembly defects that resemble those of degradation-resistant Cin8 mutants. Reduction of Cin8 SUMOylation partly stabilizes spindles in *ase1Δ* cells. In addition, SUMOylation of Cin8 and Slx5 are required for timely actomyosin-ring contraction and Cin8 clearance from the mitotic spindle. We propose that SUMOylation regulates Cin8 activity and that SUMO-targeted ubiquitylation facilitates removal Cin8 from the spindle, promoting spindle disassembly at the end of mitosis.

## INTRODUCTION

Irreversible transitions during the cell cycle are achieved by coupling the Cdk1 phospho-regulatory network to ubiquitin-dependent degradation. Spindle disassembly in late mitosis requires decrease of mitotic Cdk1 activity that promotes inactivation of factors required for spindle elongation (Sullivan and Morgan, 2007). In parallel, proteasomal degradation clears microtubule (MT) cross-linking proteins that are involved in mitotic spindle stabilization, such as the proteins Ase1/PRC1 and the yeast kinesin-5 Cin8 (Hildebrandt and Hoyt, 2001; Juang et al., 1997; Duellberg et al., 2013).

Kinesin-5 motors are conserved homotetramers essential for spindle bipolarity (Mann and Wadsworth, 2019; Pandey et al., 2021). In budding yeast, the paralogs Cin8 and Kip1 are required for spindle formation and generate forces for spindle elongation and chromosome separation during anaphase B (Saunders and Hoyt, 1992; Hoyt et al., 1992; Roof et al., 1992). Similar to other kinesin-5 motors, Cin8 crosslinks MTs (Kapitein et al., 2005; Hildebrandt et al., 2006). In metaphase, it localizes to interpolar and kinetochore MTs focusing kinetochore bundles (Gardner et al., 2008). During anaphase, Cin8 translocates to the spindle midzone, utilizing its plus-end-directed motility to elongate the spindle by sliding MT antiparallel MTs apart (Gerson-Gurwitz et al., 2011; Saunders et al., 1995; Straight et al., 1998) and stabilizing the midzone though its interaction with Ase1/PRC1 (Khmelinskii et al., 2007, 2009).

Degradation of Cin8 at the end of mitosis requires the activity of the ubiquitin ligase APC^Cdh1^ and depends on its nuclear localization signal and a bipartite destruction sequence composed of a KEN box and residues of the downstream 22 amino acids (Hildebrandt and Hoyt, 2001). Although mutations in the KEN sequence do not display a severe cell growth phenotype, Cin8 over-expession is toxic and results in spindle over-elongation (Saunders et al., 1997). Moreover, cells bearing the *cdh1Δ* deletion abolish degradation of Cin8 at the end of mitosis and display defects in a spindle disassembly that are demonstrated by over-elongated, bent spindles that acquire a fishhook shape (Woodruff et al., 2010).

Spindle over-elongation is also the phenotype of cells that are defective in removal of Cin8, Ase1 and other factors from the spindle. Three Cdk1/Cdc28 sites in the Cin8 motor domain (Avunie-Masala et al., 2011; Goldstein et al., 2017; Shapira and Gheber, 2016) differentially regulate Cin8 association on the spindle at different times during anaphase. Mutants that abolish phosphorylation at these sites display spindle disassembly defects and show persistent Cin8 localization on spindle MTs, spindle over-elongation and bending (Avunie-Masala et al., 2011).

A number of genetic approaches have identified additional pathways that are required for spindle disassembly (Woodruff et al., 2010; Vizeacoumar et al., 2010; Pigula et al., 2014). Loss-of-function mutants of the yeast Aurora B kinase Ipl1 have over-elongated spindles (Buvelot et al., 2003) since they fail to promote MT depolymerization induced by phosphorylation of MT-associated proteins at the midzone (Kotwaliwale et al., 2007; Buvelot et al., 2003). Kinesin-dependent transport of Ipl1 to the midzone is required for normal spindle disassembly (Ibarlucea-Benitez et al., 2018) and requires downregulation of Cdc28/Cdk1-dependent phosphorylation and the function of the kinetochore Ctf19 complex, mutations in which lead to formation of spindle fishhooks (Vizeacoumar et al., 2010).

Intriguingly, SUMOylation, a ubiquitin-family modification, plays an important role in Ipl1 midzone localization and spindle disassembly. Deletion of the SUMO E3 ligase Mms21 or loss of SUMOylation of the Ctf19 complex protein Mcm21 fail to localize Ipl1 to the midzone and display spindle over-elongation (Vizeacoumar et al., 2010). Similarly, SUMOylation of the kinetochore proteins Ndc10 and Cep3 is required for its localization to the spindle midzone and spindle length control (Montpetit et al., 2006).

SUMO-targeted ubiquitylation is also required for spindle disassembly. SUMO-targeted Ubiquitin Ligases are a special class of conserved ubiquitin E3 enzymes that recognize and ubiquitylate SUMOylated proteins, targeting them often for proteasomal degradation (Tatham et al., 2008; Xie et al., 2007; Uzunova et al., 2007). STUbLs contain SUMO interaction motifs (SIMs) that mediate recognition of SUMOylated substrates (Lascorz et al., 2022). Apart from a number of other functions (Chang et al., 2021; Gotthardt et al., 2025), STUbLs are implicated in quality control of kinetochore proteins and spindle-associated factors in yeast and mammalian cells (Greenlee et al., 2018; Alonso et al., 2012; Abrieu and Liakopoulos, 2019; Mukhopadhyay et al., 2010). The yeast STUbL Uls1 and the heterodimeric Slx5/Slx8 ubiquitylate the spindle positioning protein Kar9 when it mislocalizes at the plus-ends of kinetochore MTs instead of the astral MT plus-ends (Schweiggert et al., 2016b). In this case, STUbLs seem to act as QC factors that monitor correct occupancy but also proteostasis of MT-associated proteins (Schweiggert et al., 2016a; Alonso et al., 2012). Loss of STUbL function causes chromosome segregation defects in mammalian cells (Hirota et al., 2014a) and abnormally elongated fishhook spindles in yeast (van de Pasch et al., 2013).

We show here that SUMOylation and the yeast STUbL Slx5 regulate spindle disassembly via the yeast Cin8 kinesin-5. We found that Cin8 is SUMOylated and that it interacts with Slx5, suggesting that it is a STUbL substrate. Lack of Cin8 SUMOylation or of the *SLX5* gene caused phenotypes shared with cells expressing the Cin8-KED mutant that is insensitive to APC^Cdh1^-dependent degradation. Moreover, a mutant with reduced Cin8 SUMOylation was able to partly restore spindle length in *ase1Δ* deletion cells, whereas Cin8 SUMOylation and Slx5 were required for timely mitotic exit. We propose that Cin8 SUMOylation and Slx5 regulate spindle disassembly by facilitating Cin8 clearance from the spindle at the end of mitosis.

## RESULTS

### 1. Cin8 is SUMOylated *in vivo* and *in vitro*

In a proteomic screen for SUMOylation sites, we have identified Cin8 as a SUMOylated protein (Psakhye and Jentsch, 2012). SUMOylation takes place at lysine 305* (*see in Materials and Methods), which is situated in the large loop-8 of the protein that is embedded in the motor domain (Fig. 1A). Since loop-8 is one of several important regulation sites for Cin8 (Singh et al., 2018; Avunie-Masala et al., 2011), we set out to verify, whether Cin8 is indeed SUMOylated *in vivo*. After immunoprecipitation of a triple HA-tagged version of the kinesin (Cin8-HA), we detected a co-precipitating, slower-migrating Cin8-HA species that interacted with anti-Smt3 antibodies (Fig. 1B). This species was absent in the *smt3-331* mutant that has reduced SUMOylation levels at the respective temperature, suggesting that it could indeed represent a SUMO-conjugated form of Cin8.

**Figure 1:**
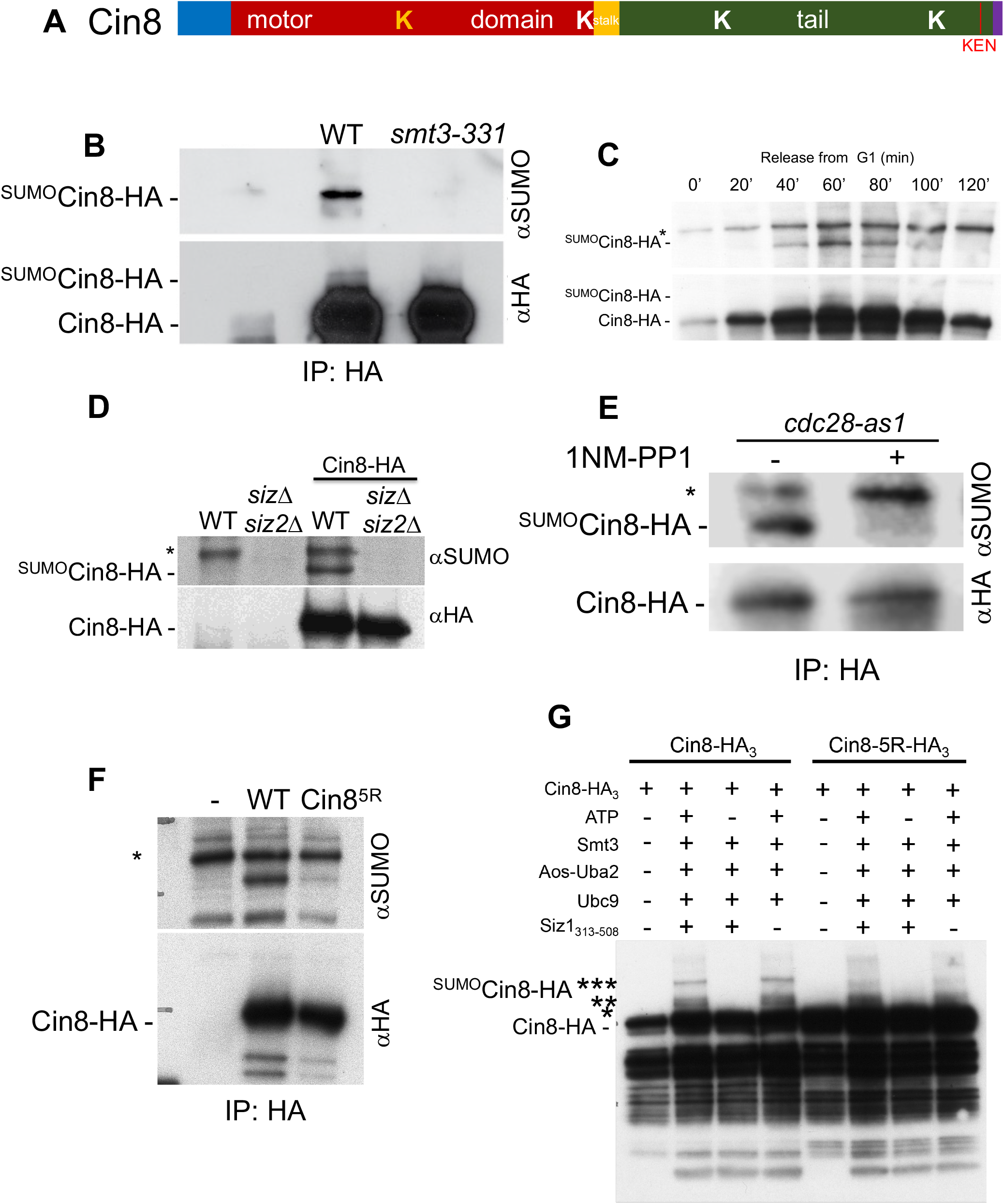
The yeast kinesin-5 Cin8 is SUMOylated *in vitro* and i*n vivo*. **A:** Scheme of the Cin8 protein with SUMOylation sites. In orange the K305 site, identified by mass spectrometry, grey: putative SUMOylation sites identified using bioinformatics. All 5 sites were substituted by arginines to generate the mutant Cin8-5R. **B:** Immunoprecipitation of Cin8-3xHA, followed by western blot using anti-SUMO antibodies reveals a Cin8 species with higher mobility likely corresponding to SUMOylated Cin8 (^SUMO^Cin8), since it is absent in the SUMO *smt3-331* mutant at the restrictive temperature. **C:** Immunoprecipitation of Cin8-3xHA from extracts of cells, after release from alpha-factor-mediated G1 arrest, followed by western blot using anti-SUMO antibody. The SUMOylated form of Cin8 follows the abundance of Cin8 during the cell cycle, (*) denotes a cross-reactive band. **D:** Inhibition of Cdc28/Cdk1 activity after treatment of *cdc28-as1* cells with the 1NP-PP1 inhibitor results in loss of Cin8 SUMOylation in immunoprecipitation/westerns as in (A) and (B) above. **E:** The SUMOylated form of Cin8 is absent in a double deletion mutant of the SUMO ligases Siz1 and Siz2, immunoprecipitation/westerns as above. **F:** The Cin8-5R-3xHA mutant has significantly reduced SUMOylation, as revealed by immunoprecipitation/anti-SUMO western blot (see above). **G:** *In vitro* SUMOylation of immunoprecipitated Cin8-3xHA and Cin8-5R-3xHA reveals a series of SUMOylated species that are less abundant in the Cin8-5R mutant. According to the molecular weights, species (*) and (**) correspond likely to mono-SUMOylated Cin8, and species (***) to di-SUMOylated Cin8. Therefore, the SUMOylated *in vivo* species in (A) corresponds likely to di-SUMOylated Cin8.

We followed the modification of Cin8 during the cell cycle by releasing cells from a G1 arrest (Fig. 1C). The modified form of Cin8 followed the amount of the protein during the cell cycle, peaking during metaphase and decreasing at the end of the cell cycle. We next examined whether the observed modification depends on the activity of the yeast SUMO ligases Siz1 and Siz2, performing analogous Cin8-HA immunoprecipitations followed by anti-Smt3 western blots. As expected, the anti-Smt3-interacting adduct was absent in the double *siz1siz2ΔΔ* mutant (Fig. 1D). To examine, whether Cdc28/Cdk1 activity is required for Cin8 modification, we used the *cdc28-1as* mutant (Bishop et al., 2000) and found that co-precipitating Smt3 modification was lost after specific inhibition of the Cdc28-1as activity (Fig.1E).

We subsequently set out to generate mutants that would abrogate SUMOylation. We mutated the K305 site, but the putative SUMOylated Cin8 species was still present in the Cin8-K305R mutant, leading us to search for other possible SUMOylation sites. After analyzing the Cin8 sequence using JASSA (Zagury et al., 2015), we identified 4 additional sites displaying high probability for SUMOylation (Fig. 1A). Sequential mutation of these sites gradually reduced Cin8 SUMOylation *in vivo* (Fig. S1A), whereas modification of the mutant Cin8-5R bearing five K->R substitutions at positions* 305, 542, 707, 854 and 1036 (Fig. S1A; *see note in Materials and Methods) was severely reduced (Fig. 1A, F). Finally, we tested SUMOylation of Cin8 and the Cin8-5R mutant *in vitro*. In the assay, we used *in vitro* purified SUMOylation enzymes and Cin8 immunoprecipitated from cells, to preserve any Cin8 posttranslational modifications (i.e. phosphorylation) possibly required for SUMOylation. We detected 3 prominent modified Cin8 species, as well as a number of higher molecular weight conjugates, which were ATP-dependent and clearly reduced in the Cin8-5R mutant. Together, these results indicate that Cin8 is likely multi-SUMOylated in a Siz1/2 and Cdk1/Cdc28-dependent manner.

### 2. Slx5 and SUMOylation of Cin8 regulate spindle length and midzone organization

Using a yeast 2-hybid assay (Y2H), we asked, whether Cin8 interacts with STUbLs. Indeed, we could detect an interaction between Slx5 and Cin8 (Fig. 2A). The interaction was still detectable with the Cin8-5R mutant, however it was absent upon mutation of the SIM motifs of Slx5 (Fig. 2A; Uzunova et al., 2007). This result suggests that Cin8 could be a substrate of Slx5.

**Figure 2:**
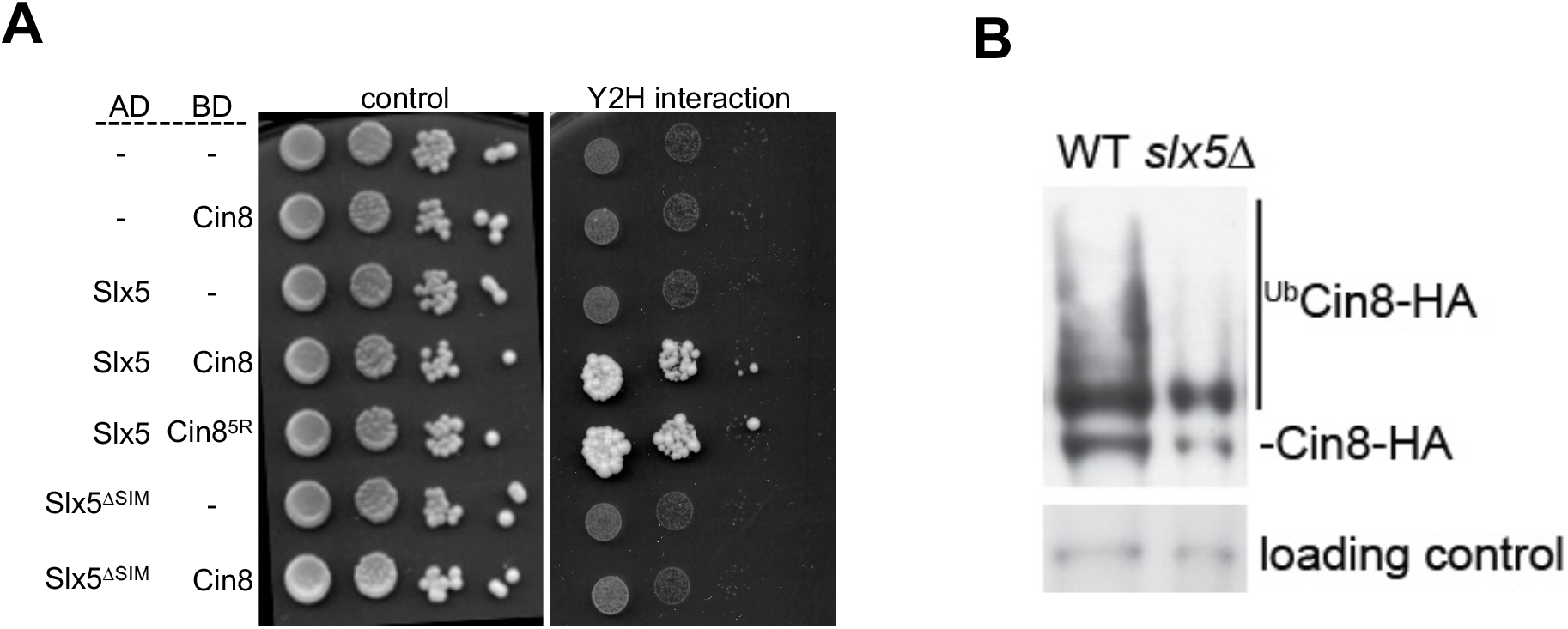
Cin8 is a substrate of Slx5. **A**: Cin8 interacts with Slx5 in a yeast two-hybrid (Y2H) assay. The Cin8-5R mutant interacts still with Slx5, but the interaction is lost upon mutation of the SIM motifs in Slx5 (Slx5^*Δ*SIM^). **B:** Ubiquitylation of Slx5 is reduced in the *slx5Δ* deletion strain. *cdc15-2 and cdc15-2 slx5Δ* cells were arrested in anaphase (inactive APC^Cdh1^), expression of His6-tagged ubiquitin was induced (+) and ubiquitylated proteins were isolated by Ni^2+^ affinity purification under denaturing conditions and analyzed for Cin8-3xHA via western blotting.

To further examine this possibility, we tested whether ubiquitylation of Cin8 is reduced in *slx5Δ* deletion mutants, using his_6_-ubiquitin pulldowns under denaturing conditions. To avoid Cin8 ubiquitylation by the APC^Cdh1^, that might mask the Slx5-dependent ubiquitylation, we performed our assays in the background of the *cdc15-2* mutation at restrictive temperature, to arrest cells in anaphase, when APC^Cdh1^ is still inactive. Indeed, we detected a reduction in Cin8 ubiquitylation in *cdc15-2 slx5Δ* cells compared to the *cdc15-2* control (Fig. 2B), suggesting that Cin8 is ubiquitylated by Slx5 *in vivo*.

We next examined the functional implications of Slx5-dependent ubiquitylation in Cin8 function. First, we analyzed the localization of Cin8-5R and Cin8 in *slx5Δ* cells during metaphase (Fig. 3A,B). Cin8 has a bi-lobed localization, concentrating to the kinetochore MTs at metaphase, and re-localizes on the spindle midzone during anaphase. In metaphase, we observed that the amount of Cin8 at kinetochores in *slx5Δ* cells was higher compared to wild-type (Fig 3A). Quantification in *slx5Δ* cells arrested at metaphase confirmed this observation and also revealed a smaller, but significant increase of Cin8-5R at kinetochores as well (Fig. 3B). This data suggested the possibility that the amount of Cin8 at spindle kinetochore MTs could be regulated by SUMO-dependent ubiquitylation.

**Figure 3:**
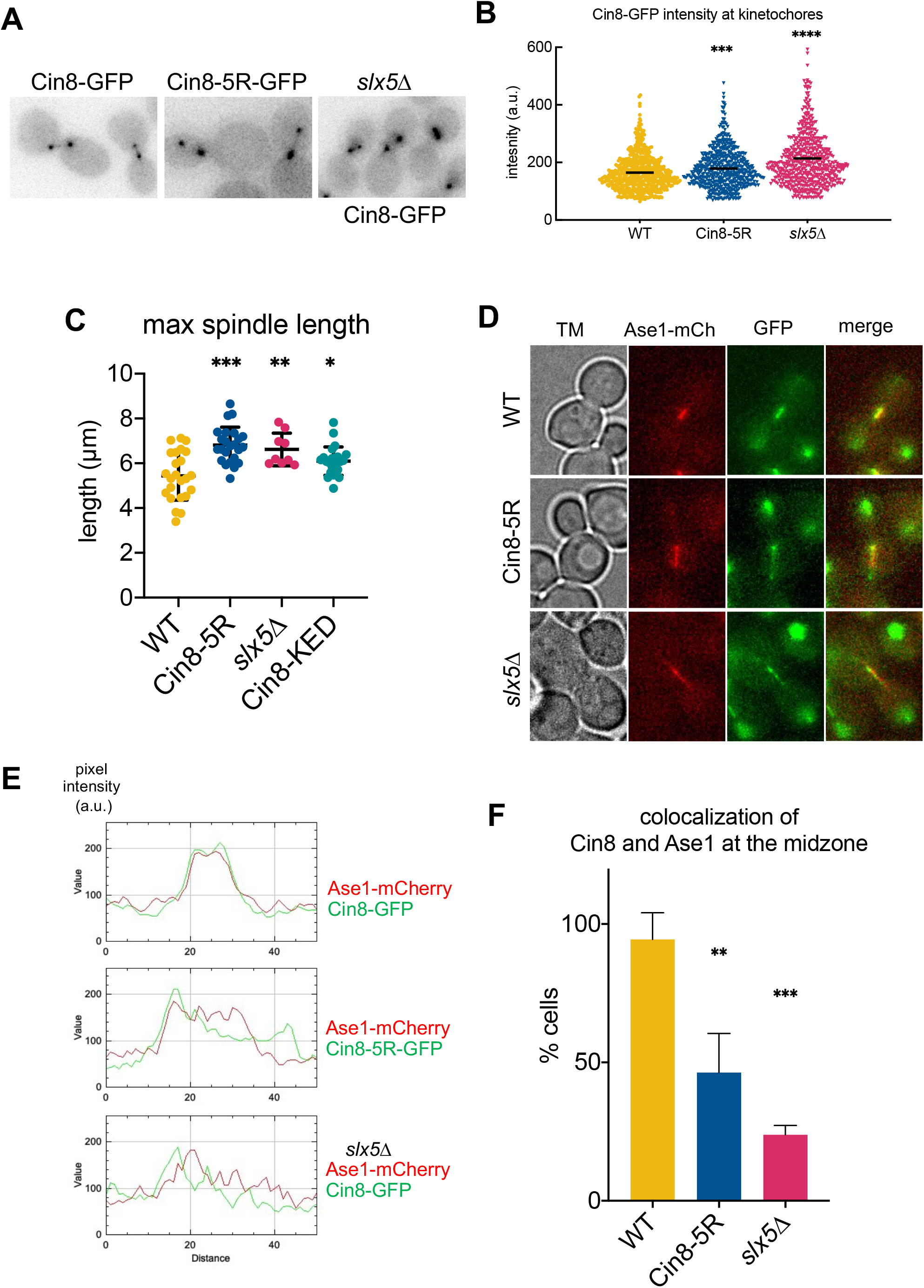
Cells lacking Slx5 and Cin8 SUMOylation display spindle defects similar to cell expressing non-degradable Cin8. **A:** Images of wild-type or *slx5Δ* cells expressing Cin8-GFP or Cin8-5R-GFP-expressing cells arrested at metaphase through Cdc20 depletion. **B:** Quantification of the GFP fluorescence in (A). The kinetochore levels of Cin8-5R-GFP or of Cin8-GFP in *slx5Δ* cells are elevated compared to wild-type, N>490, comparison: non parametric Mann-Whitney test, **p<0.01, ****p<0.0001 compared to wild-type. **C:** Cells expressing under-SUMOylated Cin8-5R or *slx5Δ* null cells display over-elongated spindles, similar to cells that express Cin8-KED that is defective in its APC-dependent degradation. Shown are measurements of maximum spindle lengths from max. intensity projections of time lapse movies of cells expressing GFP-Tub1, N=24 cells imaged for each strain, comparison: non parametric Mann-Whitney test, *p<0.1, **p<0.01, ***p<0.0001 compared to wild-type. **D:** Defects in spindle midzone organization of cells expressing under-SUMOylated Cin8-5R or *slx5Δ* null cells. The frequently perfect co-localization between Ase1-mCherry and Cin8-GFP observed in wild-type cells is perturbed in the mutants. Shown are characteristic images of cells in anaphase. **E:** Intensity plot analysis of the images in (D). Note the non-overlapping maxima of Ase1 and Cin8 in the mutants, compared to the near perfect overlap in wild-type cells. **F:** Quantification of Cin8-Ase1 co-localization in the different mutants, N>60 cells total from time lapse movies from each strain, from 3 independent experiments, comparison: non parametric Mann-Whitney test, **p<0.01, ***p<0.0001 compared to wild-type. See also Supplementary Movie S1-S6.

Second, we analyzed spindle morphology during anaphase. Cells deleted for the *SLX*5 gene display often over-elongated, fishhook anaphase spindles (van de Pasch et al., 2013). We thus asked whether cells expressing the SUMOylation-defective Cin8-5R have a similar phenotype and compared maximum anaphase spindle lengths between *slx5Δ* cells, cells expressing Cin5-5R and cells expressing the Cin8-KED mutant, that is defective in APC^Cdh1^-dependent Cin8 degradation due to a N-to-D mutation in the KEN box of the kinesin (Hildebrandt and Hoyt, 2001). Interestingly, all three mutants showed spindles that were significantly longer than wild-type (Fig. 3C). Therefore, loss of Cin8 SUMOylation and deactivation of Slx5 have similar phenotype with cells defective in Cin8 degradation. Together with the Y2H and ubiquitylation data, these results are in agreement with the hypothesis that SUMO-dependent ubiquitylation of Cin8 is required for Cin8 degradation.

We next examined the localization of Cin8 and Cin8-5R on anaphase spindles (Fig. 3D-F, Supplementary Movie S1-S6). In cells with elongated anaphase spindles, Cin8-5R-GFP was often mislocalized, either along the entire spindle or forming “patches” and failing to properly focus at the midzone. To better characterize the phenotype, we used time-lapse microscopy and followed anaphases of cells co-expressing the midzone marker Ase1-mCherry, quantifying the percentage of cells that failed to perfectly co-localize the proteins at the midzone (Fig. 3F). Indeed, more than 45% of Cin8-5R-expressing cells displayed partial or complete mislocalization at the Ase1 midzone, in contrast to over 95% of wild-type cells displaying perfect colocalization of the 2 proteins. We observed the same phenotype in *slx5Δ* cells, with over 70% of the population failing to perfectly co-localize. These results are in agreement with previous findings showing that increasing the amount of Cin8 in cells (due to lack of degradation) or on the spindle (due to loss o Cin8 phosphorylation) leads to spindle over-elongation (Avunie-Masala et al., 2011; Woodruff et al., 2010). They also suggest that SUMOylation and SUMO-targeted ubiquitylation is required to control the amount and distribution of Cin8 on the mitotic spindle, both in metaphase and anaphase cells.

### 3. Lack of Cin8 SUMOylation partly recues the phenotype of *ase1Δ* cells

The results obtained until now showed that downregulation of Cin8 SUMOylation and deletion of *SLX5* result in phenotypes that are similar to the phenotypes of cells expressing the non-degradable Cin8-KED. To further investigate this finding, we asked whether Cin8-5R could suppress the short spindle phenotype of *ase1Δ* cells since Ase1 and Cin8 form a functional unit and Ase1 is required to recruit Cin8 to the spindle (Khmelinskii et al., 2009). Therefore, Cin8 “gain of function” may rescue the phenotype of *ase1Δ* cells. We thus used time-lapse microscopy to follow anaphases of *ase1Δ* cells expressing mCherry-TUB1 (α-tubulin) and Cin8-GFP or Cin8-5R-GFP (Fig. 4A). Spindles expressing Cin8 in absence of the *ASE1* gene broke often and resumed elongation, remaining always shorter compared to wild-type, as already described ((Schuyler et al., 2003), Fig. 4A,C). Spindles of cells expressing Cin8-5R in the presence of the *ASE1* gene spent more time in anaphase and elongated excessively before disassembly, consistent with our prior findings (Fig. 4A). Interestingly, spindles of *ase1Δ* Cin8-5R cells grew longer compared to *ase1Δ* Cin8-GFP spindles at all time points (Fig. 4A, C) and displayed a significantly greater maximum length compared to those of Cin8 expressing *ase1Δ* cells (Fig. 4B). Hence, the Cin8-5R mutant can partly suppress the short spindle phenotype of *ase1Δ* cells. Intriguingly however, expression of Cin8-KED did not have the same effect. Spindles of *ase1Δ* Cin8-KED cells displayed lengths similar to the spindles of *ase1Δ* Cin8 cells (Fig. S1B), suggesting that the Cin8-5R mutant is not always equivalent to Cin8-KED. Thus, these experiments are in line with the notion that loss of Cin8 SUMOylation leads to some “gain of function” of the kinesin.

**Figure 4:**
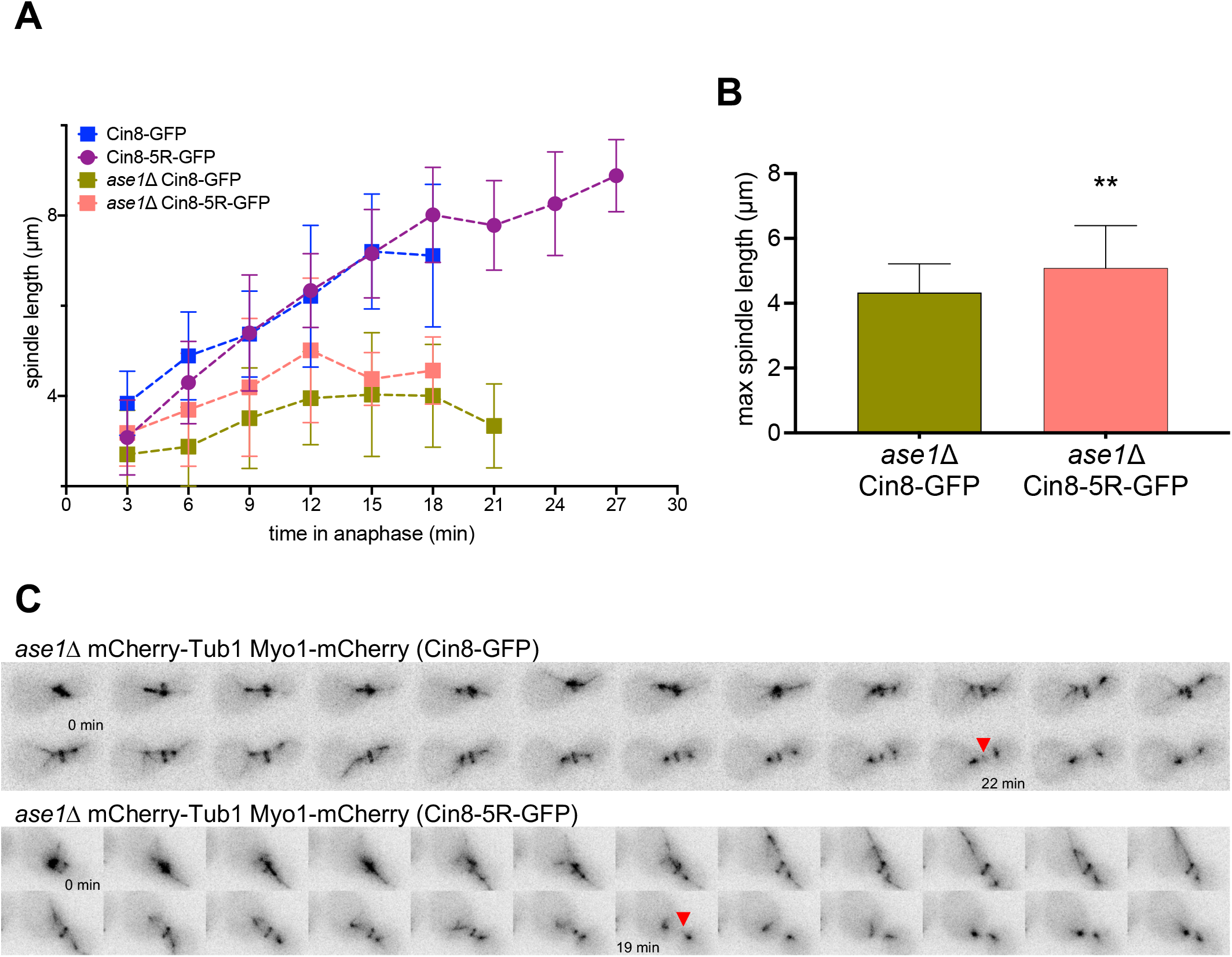
The Cin8-5R mutant partly rescues the phenotype of *ase1Δ* cells. **A:** Measurements of spindle lengths during anaphase spindle elongation show that spindles in *ase1Δ* Cin8-5R cells are constitutively longer than the spindles of *ase1Δ* Cin8-GFP cells. Spindles of Cin8-5R-expressing cells not deleted for the *ASE1* gene keep extending for a longer time, resulting in spindle over-elongation, see also Fig. 3C. Plotted are mean values +/-SD, N>13 cells followed in time-lapse movies for each strain. **B:** The maximum spindle length achieved by Cin8-5R-expressing *ase1Δ* cells in greater compared to *ase1Δ* Cin8-GFP cells, N>13 cells followed in time-lapse movies for each strain, comparison Welch’s t test, **p<0.01 **C:** Example of *ase1Δ* cells in (B) expressing mCherry-Tub1, Myo1-mCherry and Cin8-GFP or Cin8-5R-GFP, performing anaphase spindle elongation (1 min interval between frames). Note that the spindles grow longer in *ase1Δ* Cin8-5R-GFP cells, and that the spindle MT density is very low in *ase1Δ* Cin8-GFP cells. In addition, the end of cytokinesis (red arrowhead) occurs later in the *ase1Δ* Cin8-GFP cell compared to *ase1Δ* cells expressing Cin8-5R-GFP, see also Fig. 5C. The GFP channel is now shown (brackets).

### 4. Cin8 SUMOylation is required for timely spindle disassembly at the end of mitosis

The experiments until know were consistent with the requirement of SUMOylation and subsequent Slx5-dependent ubiquitylation for degradation or otherwise deactivation of the Cin8 kinesin. We subsequently asked whether SUMO/Slx5 is required for removal of Cin8 from the anaphase spindle. Using time-lapse microscopy of mCherry-Tub1 Myo1-mCherry Cin8-GFP expressing cells, we compared the duration between the beginning of spindle elongation and the complete clearance of Cin8-GFP from the spindle between the wild-type, the *slx5Δ* mutant and cells expressing different versions of Cin8 (Fig. 5A-C). In addition, we measured the duration between anaphase entry and the disassembly of the actomyosin ring (AMR) using respectively, the mCherry-Tub1 and the Myo1-mCherry signal as a reference (Fig. 5A-C, See also Supplementary Movie S7-17).

**Figure 5:**
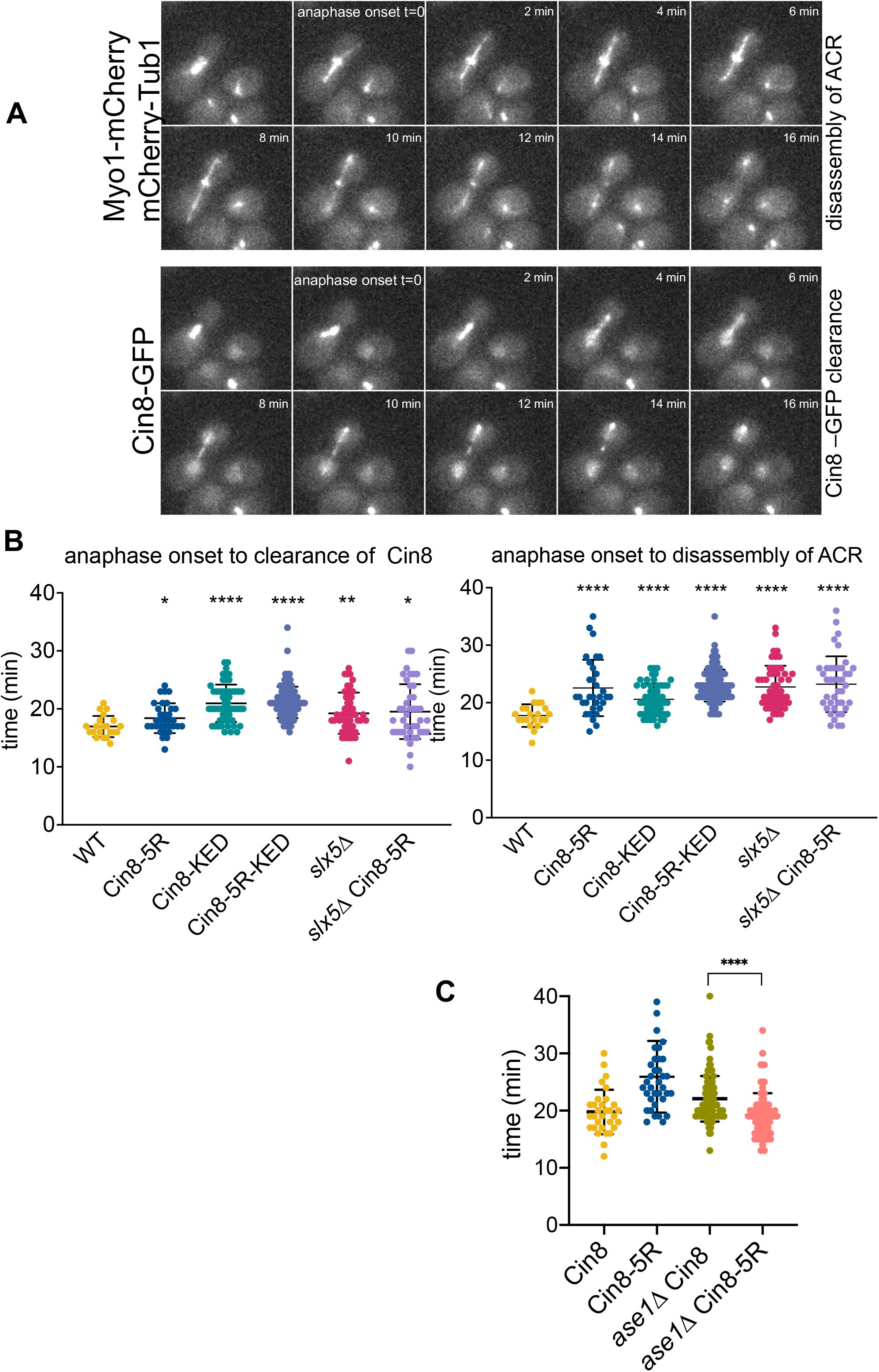
Slx5 and Cin8 SUMOylation is required for timely anaphase. **A:** Example of Cin8-GFP Myo1-mCherry cell performing anaphase, illustrating the measurements shown in B and C below. The duration of 2 events was measured a) the time between start of spindle elongation and clearance of Cin8 from the spindle b) the time between start of spindle elongation and complete disassembly of AMR. **B: (left)** The non-degradable Cin8-KED mutant displays delayed clearance of Cin8 from the spindle. The same holds true for Cin8-5R-GFP and *slx5Δ* Cin8-GFP cells. (**right**) All mutants display a delay regarding the timing between anaphase onset and the disassembly of the AMR, comparison: non-parametric Mann-Whitney test, *p<0.1, **p<0.01, ***p<0.0001, ****p<0.0001. Duration of events in Fig. S1C. See also Supplementary Movie S7-17 **C:** Cells deleted for *ase1Δ* also display a delay in the timing between anaphase onset and the disassembly of the AMR. Expression of Cin8-5R-GFP restores this delay to wild-type levels, see also Fig. 4C, comparison: non-parametric Mann-Whitney test, ****p<0.0001. Duration of events in Fig. S1D.

All single mutants displayed significant and reproducible delays in clearance of Cin8 from the spindle (Fig. 5B, C). The delays were of the same order, with the minimum of 1,4 min for Cin8-5R and a maximum 4,0 min for Cin8-KED. The Cin8-5R-KED double mutant, bearing both 5R and KED mutations, behaved essentially as Cin8-KED, showing a clearance delay of 4,1 min. The *slx5Δ* cells showed an intermediate delay of app. 2,5 min, whereas expressing Cin8-5R in *slx5Δ* cells showed essentially the same timing of Cin8 clearance.

In addition to the clearance phenotype, both Cin8-KED cells and STUbL pathway mutants displayed delays in AMR contraction (Fig. 5B,C). Here, Cin8-KED had the smallest delay of 2,8 min, whereas Cin8-5R disassembled the AMR 4,8 min later than wild-type, that corresponds to an increase of 27% compared to wild-type. The Cin8-5R-KED mutant behaved similar to Cin8-5R delaying AMR disassembly by 5,2 min. Finally, *slx5Δ* cells displayed a similar delay of app. 5 min in AMR contraction, irrespective of whether they expressed Cin8 or Cin8-5R.

We also tested anaphase timing in *ase1Δ* cells (Fig. 5C). Intriguingly, (Fig. 5A) *ase1Δ* Cin8 cells displayed a delay in the timing of AMR contraction relative to wild-type. This delay was however restored to wild-type values, when Cin8-5R was expressed instead of Cin8.

This is consistent with the previous data on spindle lengths (Fig. 4A, B) showing that expression of Cin8-5R suppresses the defects of *ase1Δ* cells, at least partly.

In conclusion, these observations add to the findings suggesting that abrogation of Cin8 SUMOylation or deletion of the *SLX5* gene result in phenotypes similar to those caused by failure to degrade Cin8 due to the KED mutation. They are therefore consistent with the notion that SUMO-dependent ubiquitylation is required for degradation and removal of Cin8 from the mitotic spindle.

## DISCUSSION

STUbLs control chromosome segregation and spindle dynamics ((Hirota et al., 2014b; van de Pasch et al., 2013). Using Y2H assays and mass spectrometry to identify SUMOylation sites, we have identified the yeast Cin8/kinesin-5 as a Slx5 substrate and could demonstrate that Cin8 is SUMOylated *in vivo* and *in vitro*. Interestingly, the SUMOylation site K305 resides in the loop-8, an important site for Cin8 binding to MTs and regulation of Cin8 clustering and directionality (Gerson-Gurwitz et al., 2011). We could clearly detect at least 2 SUMOylated species, and needed to mutate 5 SUMOylation sites in total, to achieve reduction in SUMOylation. Modification of Cin8 by SUMO depends on the SUMO ligases Siz1/2, follows the amount of the kinesin during the cell cycle and peaks with maximum levels of Cin8. In addition, it requires Cdk1/Cdc28 activity. Together, these data suggest that SUMOylation of Cin8 exerts its function during mitosis.

Abrogation of SUMOylation does not reduce the interaction of Cin8 with Slx5 in Y2H. This is common to other STUbL substrates, i.e. □ □ □α□ and Kar9 (Schweiggert et al., 2016b; Xie et al., 2010). SUMO is not the only determinant of the interaction between STUbLs and their substrates (Hickey and Hochstrasser, 2015). SUMO-SIM interactions are low affinity and group SUMOylation often provides a platform for recruitment of STUbLs, making the interaction insensitive to the mutation of SUMOylation sites of only one component. We suspect that other Cin8-interacting spindle proteins are SUMOylated, as is the case for Ase1 (our unpublished data). The fact that the interaction between Slx5 and Cin8 is SIM-dependent suggests that Cin8-STUbL interaction may be occurring in the context of a group SUMOylated complex.

Over-elongated spindles are a feature of spindle disassembly defects, both in meiosis II and mitosis (Woodruff et al., 2010; Seitz et al., 2023). They are also a phenotype of Cin8 *cdh1Δ* cells or of cells that have difficulty to remove Cin8 from the spindle: cells expressing non-phosphorylatable Cin8-AAA mutants that remain attached to the spindle (Avunie-Masala et al., 2011) display spindle over-elongation and spindle bending. Our results show that cells expressing Cin8-KED, that cannot be degraded by APC^Cdh1^, also display spindle over-elongation (Fig. 3C) and delayed clearance of Cin8 from the spindle (Fig. 5B), indicating at least part of the *cdh1Δ* spindle disassembly defects can be attributed to increased Cin8 activity due to failure to degrade Cin8 at the end of mitosis.

Importantly, both Cin8-5R and *slx5Δ* behave similar to the non-degradable Cin8-KED mutant in most cases: they have over-elongated spindles, display delayed clearance of Cin8 from the spindle at telophase and delay the onset of AMR contraction (Fig. 5B). These phenotypes are consistent with increased amounts or activity of Cin8 on the spindle in these mutants. However, Cin8-5R partly recues the spindle phenotype of *ase1Δ* cells, resulting in some spindle stabilization, but the Cin8-KED mutant does not have this effect. The reason for this difference is not clear. SUMOylation may regulate the activity of Cin8 in a degradation-independent manner, in addition to its role in the Slx5-dependent degradation of the kinesin. Nevertheless, most phenotypic data suggest, together with the Y2H Cin8-Slx5 interaction, the reduction of Cin8 ubiquitylation in *slx5Δ* mutants and the increased localization at kinetochores in *slx5Δ* and Cin8-5R-expressing cells (Fig. 3A), that Slx5 facilitates spindle disassembly by down-regulating Cin8. This is also in line with the negative synthetic genetic interactions of *slx5Δ* with mutants in genes involved in spindle disassembly pathways: *mcm21, dcc1, cdh1, ctf8* and *ipl1* ((Woodruff et al., 2010)).

Intriguingly, we found that Cin8-5R-expressing, *slx5Δ* but also *ase1Δ* cells have a significant and reproducible delay in the time required from spindle elongation until the disassembly of the AMR. In contrast, constriction of the AMR occurred with normal kinetics in all mutants tested. The cause of this delay is not clear. Since all these mutants perturb anaphase spindle dynamics and structure, this finding points to a role of the spindle in regulating activation of FEAR or MEN and the timing of mitotic exit.

The cellular pool of Cin8 is degraded at the end of mitosis by APC^Cdh1^-dependent degradation. We favor a model in which Slx5 ubiquitylates SUMOylated Cin8 resulting in its removal from the spindle at the end of mitosis. What is the use of a second STUbL-dependent pathway? The APC cannot access proteins in residing in MT bundles (Song et al., 2014), like the ones at kinetochores or the midzone, but can degrade the bulk of MT-unbound, nucleoplasmic Cin8. Therefore, modification of SUMO and extraction by STUbLs may facilitate APC-dependent degradation. Moreover, the fact that combining the 5R and the KED mutations does not exacerbate any phenotype suggests that STUbL and APC-dependent degradation may be linked. In agreement with these ideas, a number of midzone proteins including the centralspindlin MKLP-1/Kif23 are heavily SUMOylated (Abrieu and Liakopoulos, 2019).

The mechanism of STUbL-dependent Cin8 spindle removal could be complementary to phosphorylation, since Cdk1/Cdc28-dependent phosphorylation is able to detach kinesin-5 from the spindle (Shapira and Gheber, 2016). However, at late mitosis the activity of Cdk1 is reduced, and dephosphorylation of Cin8 by the PP2A and Cdc14 phosphatases results in Cin8 accumulation near the spindle poles and at the spindle midzone (Roccuzzo et al., 2015; Avunie-Masala et al., 2011). Therefore, STUbL-dependent ubiquitylation could help to mobilize/clear Cin8 from the spindle at this stage.

Another possibility that does not exclude the previous hypotheses is that Slx5 acts as quality control factor (Guo et al., 2014; Keiten-Schmitz et al., 2020; Ohkuni et al., 2016; Folger et al., 2025). In the case of Kar9, localization of the protein and its amount on MT are regulated (Schweiggert et al., 2016b). It is not clear what type of QC regulation Cin8 could be subjected to and why. One possibility is that SUMO-targeted ubiquitylation controls the amount of Cin8 on the spindle. Cin8-5R and Cin8 in *slx5Δ* cells display disrupted midzone localization, as shown by their reduced colocalization with Ase1 (Fig. 3F). Therefore, the spindle over-elongation phenotypes of these cells are likely due to perturbation of the distribution of Cin8 on the spindle. Ase1 acts as a brake for Cin8-dependent antiparallel MT sliding and spindle elongation (Braun et al., 2011). Thus, increased Cin8 amounts could disturb the stoichiometry of Cin8/Ase1 forming complexes, leading to spindle over-elongation. In this case, Slx5 would regulate the amount of Cin8 on the spindle guaranteeing correct Ase1/Cin8 stoichiometry. Lastly, it will be also interesting to investigate the importance of STUbL-dependent Cin8 degradation for subsequent cell cycles, since spindle/midzone remnants of the previous cell cycle seem to perturb spindle formation in subsequent cell divisions (Woodruff et al., 2012).

## MATERIALS AND METHODS

* There is a difference in the translational start of Cin8 between the annotated Cin8 protein in the SGD database, and the publication of (Hoyt et al., 1992) with the SGD protein being 38 aminoacids shorter than the published sequence. In the paper, we follow the original publication and thus lysines 305, 542, 707, 854 and 1036 correspond to lysines 267, 504, 669, 816, 998 in the SGD protein sequence. Irrespective of the nomenclature, all Cin8 mutants are inserted in the wild-type locus and preserve the wild-type translational initiation site.

### Yeast and Bacterial Growth Conditions, Strains, Plasmids and Primers

All strains were derivates of W303 (*leu2-3 leu2-112 trp1-1 ura3-1 his3-11 his3-15 ade2-1 can1-100*) except for the Y2H strain, which was PJ69-4A (James et al., 1996). Yeast strains are listed in Supplementary Table S1, plasmids in Supplementary Table S2 and primers in Supplementary Table S3. Media and genetic manipulations were done as in (Guthrie and Fink, 1991). The Cin8 mutations were introduced through one-step gene replacement in the Cin8 locus, leaving the promoter and translational initiation region intact. Y2H was performed as described previously (Kammerer et al., 2010). Standard molecular biology methods were used (Ausubel et al., 1989) and chromosomal deletions of open reading frames and tag insertions were performed as described (Longtine et al., 1998; Janke et al., 2004). Point mutations (for creation of the Cin8-5R and the Cin8-KED mutants) were introduced by site-directed mutagenesis (Phusion™ polymerase, Thermo Scientific).

### Biochemistry

#### Ubiquitin pulldown

experiments were carried out as described previously (Kammerer et al., 2010). For pulldowns, expression of His6-ubiquitin was induced with 1 mM CuSO_4_ for 4 h.

#### Immunoprecipitations of Cin8-3xHA and Cin8-3xHA-PrSc-MBP

Cells were grown in full YPD media until an OD 1, pelleted and lysed in 50 mM Tris pH 7.6, 150 mM NaCl, 1 mM MgCl_2,_ protease- and phosphatase-inhibitors and 0,1% NP-40. Proteins were precipitated either using 2 µg anti HA-antibody 7HA and Dynabeads™ protein A (Invitrogen), followed by elution through boiling in 2xLaemmli buffer (Cin8-3xHA), or using amylose resin (NEB) followed by elution through cleavage with PreScission protease.

#### In vitro SUMOylation reactions

were essentially performed as described (Xie et al., 2007; Johnson and Gupta, 2001) in a buffer containing 20 mM TrisCl pH7.4, 50 mM KCl, 1 mM MgCl2, 5 µM ZnCl, 5 mM ATP and 0.1 mM beta-mercaptoethanol.

#### Western blots

Protein extraction was performed as in (Knop et al., 1999) or using glass beads in 50 mM Tris pH 7.6, 150 mM NaCl, 1 mM MgCl_2,_ protease- and phosphatase-inhibitors and 0,1% NP-40. Cell lysates were analyzed by SDS-PAGE on 6% polyacrylamide gels and western blot.

#### Antibodies

monoclonal mouse α-HA (12CA-5), or clone 7HA, Sigma-Aldrich, St.-Louis, USA), α-Arc1 (polyclonal, rabbit; gift from E. Hurt (Simos et al., 1996), polyclonal rabbit α -Smt3 (Gift from F. Melchior and A. Pichler).

### Live cell imaging and analysis

Cells were grown overnight in YPD liquid cultures containing additional Adenin, Tryptophan and Uracil and imaged in a glass-bottomed Fluorodish™ (WPI) pretreated with Concanavalin A (0.5 %, 10 min), covered with an 1% agarose pad in non-fluorescent yeast medium (Waddle et al., 1996). Cells were imaged with a Nikon Ti2 inverted microscope run by the NIS Elements software, equipped by a CMOS back-illuminated Prime95B Photometrics camera (11µm pixel size), a xxx stage and a xxx Piezo, using an oil 100X Plan Apochromat 1.40 NA objective and LED light sources. Imaging was performed in a temperature-controlled chamber, at 30°C. z-stacks of 7 slices (0.4μm step/slice), from 3 or more different positions were collected. The maximum intensity projection of the z-stacked images was used to visualize Cin8-GFP and Ase1-mCherry and Myo1-mCherry or measure intensities.

*Image analysis* was done using the FIJI software. Quantification of Cin8-GFP at kinetochores was performed using “spots in yeasts“(https://napari-hub.org/plugins/spots-in-yeasts), an AI-based Napari plugin segmenting yeast cells and fluorescent spots to extract statistics, developed in collaboration with the MRI imaging facility (Clément H. Benedetti). Statistical analysis was performed with the Graphpad Prism 8 software. Samples were first tested for normality, followed by appropriate statistical testing. The number N of cells is indicated in the figure legends.

## Supporting information

Table S1 Yeast Strains

Table S2 Plasmids

Table S3 Primers

## AUTHOR CONTRIBUTIONS

F.G., D.P., D.P. I.P. and D.L. designed the experiments. F.G., D.P and I.P. carried out the experiments. F.G., D.P., D.P. and D.L. analyzed the data. D.L. wrote the manuscript. D.P., and D.L. initiated the research and supervised the work. All authors discussed the results and commented on the manuscript.

## ACKNOWLEDGEMENTS

We are grateful to late S. Jentsch and to I. Psakhye for sharing data, to L. Gheber and S. Piatti for sharing plasmids and yeast strains and to F. Melchior, J. Schweiggert and A. Pichler for the anti-Smt3 antibodies. We acknowledge the imaging facility MRI, member of the national infrastructure France-BioImaging infrastructure supported by the French National Research Agency (ANR-10-INBS-04, «Investments for the future» and the AAP ITMO INSERM 2016: Plan Cancer, (acquisition d’Equipement pour la recherche en cancérologie). D. Portran and D. Liakopoulos acknowledge the support of the French Agence Nationale de la Recherche, grant ANR-19-CE13-0012-01 (SUMOzone) and the support of the Ligue contre le cancer, contract Nr 269213. D. Panigada was supported by EMBO Long Term fellowship ALTF 911-2014.

