## Supplementary material for "Control of mitotic spindle disassembly through SUMO-targeted ubiquitylation of yeast kinesin-5": Figure S1

A

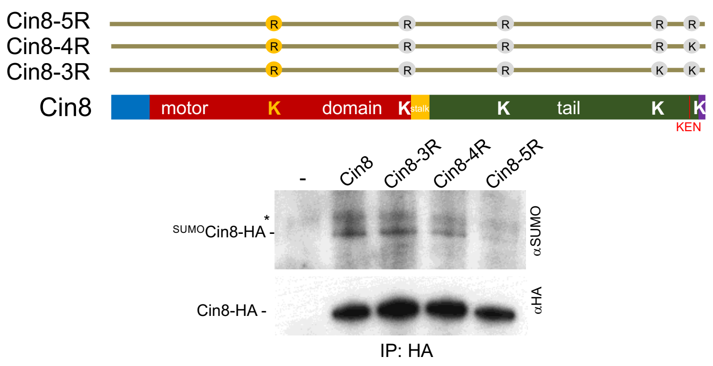

B

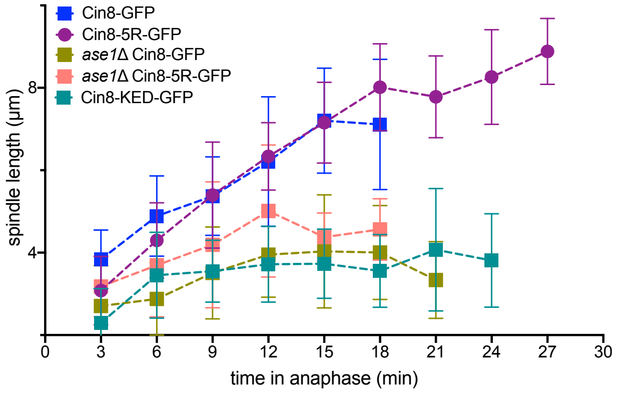

C

| | anaphase onset to clearnace of Cin8 | | | anaphase onset to disassembly of AMR | | |
|---|---|---|---|---|---|---|
| | Mean | SD | N | Mean | SD | N |
| WT | 17.0 | 1.8 | 21 | 17.8 | 2.0 | 21 |
| Cin8-5R | 18.4 | 2.6 | 33 | 22.6 | 4.9 | 32 |
| Cin8-KED | 20.9 | 3.2 | 55 | 20.6 | 2.7 | 55 |
| Cin8-5R-KED | 21.1 | 2.7 | 105 | 23.0 | 2.8 | 105 |
| *slx5*Δ | 19.2 | 3.6 | 51 | 22.7 | 3.7 | 51 |
| *slx5*Δ Cin8-5R | 19.5 | 4.7 | 41 | 23.2 | 4.8 | 41 |

D

| | Cin8 | Cin8-5R | *ase1*Δ Cin8 | *ase1*Δ Cin8-5R |
|---|---|---|---|---|
| Number of values | 31 | 36 | 104 | 78 |
| Mean | 19.8 | 25.9 | 22.1 | 19.1 |
| Std. Deviation | 3.88 | 6.27 | 3.99 | 3.96 |
| Std. Error of Mean | 0.697 | 1.05 | 0.391 | 0.449 |
