## Supplemental Information for "Control of mitotic spindle disassembly through SUMO-targeted ubiquitylation of yeast kinesin-5"

Supplementary Figure S1, Movie S1-S17, Table S1, S2, S3

**Figure S1:**

**A:** top: scheme of the constructs showing substituted Cin8 SUMOylation sites tested bottom: gradual loss of Cin8 SUMOylation *in vivo* upon sequential substitution of predicted SUMOylation sites (grey) in addition to the mass-spec identified site (orange). Shown are immunoprecipitations of Cin8-3xHA, followed by western blot using anti-SUMO antibodies. **B:** Measurements of spindle lengths during anaphase spindle elongation as in Fig. 4A including Cin8-KED, showing that spindles in *ase1*Δ Cin8-KED cells are have spindle lengths comparable to *ase1*Δ Cin8-GFP cells. Plotted are mean values +/- SD, N>13 cells followed in time-lapse movies for each strain (N=22 for Cin8-KED) **C, D:** Tables summarizing the data for Fig. 5B-C.

**Movie S1-S6:**

Movies of C-terminally GFP-tagged versions of Cin8 and Ase1-mCherry co-expressing cells in anaphase.

**Movie S7-S17:**

Movies of Myo1-mCherry mCherry-Tub1 cells expressing C-terminally GFP-tagged versions of Cin8 from metaphase until contraction of the AMR.
