## Supplementary material for "Control of mitotic spindle disassembly through SUMO-targeted ubiquitylation of yeast kinesin-5": Table S1 Yeast Strains

| Strain | Genotype | Source |
|---|---|---|
| YDL3646 | *W303 MATa*<br>*ade2-1 trp1-1 leu2,3-112 his3-11,15 ura3 can1-100 GAL+ psi+* | K699 W303 received from Simonetta Piatti |
| YDL637 | *DF5 MATa*<br>*ura3-52 lys2-801 his3-Δ200 trp1-1 leu2-3,2* | (Ulrich, 2000) |
| YDL4374 | *W303 MATa*<br>*CIN8-GFP:LEU2 mCherry-TUB1:URA3 MYO1-mCherry:hphNT1* | This Study |
| YDL4483 | *ade2-1 trp1-1 leu2,3-112 his3-11,15 ura3 can1-100 GAL+ psi+*<br>*CIN8-5R-GFP:LEU2 mCherry-TUB1:URA3 MYO1-mCherry:hphNT1* | This Study |
| YDL4563 | *W303 MATa*<br>*ase1::HIS3 mCherry-TUB1:URA3 MYO1-mCherry:hphNT1* | This Study |
| YDL4564 | *W303 MATa*<br>*ase1::HIS3 CIN8-5R-GFP:LEU2 mCherry-TUB1:URA3 MYO1-mCherry::hphNT1* | This Study |
| YDL4573 | *W303 MATa*<br>*ASE1::HIS3 CIN8-GFP:LEU2 mCherry-TUB1:URA3 MYO1-mCherry:hphNT1* | This Study |
| YDL4676 | *W303 MATa*<br>*ase1::HIS3 CIN8-KED-GFP:LEU2 mCherry-TUB1:URA3 MYO1-mCherry:hphNT1* | This Study |
| YDL4298 | *W303 MATa*<br>*CIN8-GFP:LEU2 MYO1-mCherry:hphNT1* | This Study |
| YDL4546 | *W303 MATa*<br>*CIN8-KED-GFP:LEU2 MYO1-mCherry:hphNT1* | This Study |
| YDL4551 | *W303 MATa*<br>*CIN8-5R-GFP:LEU2 MYO1-mCherry:hphNT1* | This Study |
| YDL4583 | *W303 MATa*<br>*CIN8-5R- KED-GFP:LEU2 MYO1-mCherry:hphNT1 ADE2* | This Study |
| YDL4619 | *W303 MATa*<br>*CIN8-GFP:LEU2 slx5::TRP1 MYO1-mCherry:hphNT1* | This Study |
| YDL 4645 | *W303 MATa*<br>*CIN8-5R-GFP:LEU2 SLX5::TRP1 MYO1*-mCherry:*hphNT1* | This Study |
| YDL 4715 | *W303 MATa*<br>*GAL1p-CDC20:URA3 CIN8-GFP:LEU2 MYO1-mCherry:hphNT1 ADE2* | This Study |
| YDL 4687 | *W303 MATa*<br>*GAL1p-CDC20:URA3 CIN8-GFP:LEU2 slx5::TRP1 MYO1-mCherry:hphNT1 ADE2* | This Study |
| YDL4685 | *W303 MATa*<br>*GAL1p-CDC20:URA3 CIN8-5R-GFP:LEU2 MYO1-mCherry:hphNT1 ADE2* | This Study |
| YDL4430 | *W303 MATalpha*<br>*CIN8-5R-GFP:LEU2 ASE1-mCherry:HIS3* | This Study |
| YDL 4432 | *W303 MATa*<br>*CIN8-GFP:LEU2 ASE1-mCherry:HIS3* | This Study |
| YDL 4438 | *W303 MATa*<br>*CIN8-GFP:LEU2 ASE1-mCherry:HIS3 slx5::TRP1* | This Study |

| | | |
|---|---|---|
| YDL 4553 | *W303 MATa*<br>*CIN8-KED-GFP:HIS3 ASE1-mCherry:URA3* | This Study |
| YDL3996 | *W303 MATa*<br>*CIN8-3HA:HIS3* | This Study |
| YDL3997 | *W303 MATa*<br>*CIN8-3HA:TRP1* | This Study |
| YDL3998 | *W303 MATa*<br>*CIN8-3HA:LEU2* | This Study |
| YDL4240 | S288c MATa<br>*CIN8-5R*-3HA:*LEU2* | This Study |
| YDL 4002 | W303 MATa<br>*cdc15-2 Cin8-3HA:HIS3* | This Study |
| YDL 4019 | W303 MATa<br>*cdc15-2 slx5::TRP1 Cin8-3HA:HIS3* | This Study |
| YDL4427 | W303 MATa<br>*Cin8-3HA-Precision-MBP:LEU2* | This Study |
| YDL4440 | W303 MATa<br>Cin8-5R-3HA-Precision-MBP:LEU2 | This Study |
| YDL3931 | *cin8::KanMx4 gal4Δ gal80Δ GAL2-ADE2 LYS::GAL1-HIS3*<br>*met2::GAL7-lacZ* | This Study |
| YDL4182 | *DF5 ura3-52 lys2-801 his3-Δ200 trp1-1 leu2-3,2*<br>*cdc48-6 CIN8-GFP:LEU2* | This Study |
| YDL4027 | *DF5 ura3-52 lys2-801 his3-Δ200 trp1-1 leu2-3,2*<br>*cdc48-6 GFP-TUB1-URA3* | This Study |
| YDL4141 | *S288C* MATa<br>*MET25p-CDC20-HA Cin8-HA:LEU2* | This Study |
| YDL3377 | *S288C* MATa<br>*MET25p-CDC20-HA* | This Study |
