## Supplementary material for "Control of mitotic spindle disassembly through SUMO-targeted ubiquitylation of yeast kinesin-5": Table S2 Plasmids

| Plasmid | Relevant name | Backbone | Source |
|---|---|---|---|
| pDL284 | pGAD-C1 | | (James et al., 1996)(James et al., 1996) |
| pDL285 | pGBD-C1 | | |
| pDL823 | pGAL4-AD-Slx5 | | (Schweiggert et al., 2016) |
| pDL960 | pGAL4-AD-Slx5-SIM1234 | | (Schweiggert et al., 2016) |
| pDL966 | pGBD-CIN8 | | This study |
| pDL1061 | pRS305-CIN8-Cter-3HA | pRS305 | This study |
| pDL1063 | pRS303-CIN8-Cter HA | pRS303 | This study |
| pDL1065 | pRS303-CIN8-Cter GFP | | This study |
| pDL1067 | pRS305-CIN8-Cter-GFP | pRS305 | This study |
| pDL1068 | pRS303-ASE1-Cter-mCherry | pRS303 | This study |
| pDL1166 | pRS305-CIN8Ct-3HA-PrSc-MBP | pRS305 | This study |
| pDL1168 | pRS305-CIN8-5R-Ct-3HA-PrSc-MBP | | This study |
| pDL1171 | pGBD-CIN8-5R | | This study |
| pDL1099 | pRS305-Cin8-K305R K542R K707R K854R K1036R-Cter-3HA (pRS305-Cin8-5R-Cter-3HA) | pRS305 | This study |
| pDL1176 | pRS305-Cin8-Cter-KED-GFP | pRS305 | This study |
| pDL1267 | pRS305-Cin8-5R-Cter-GFP | pRS305 | This study |
