## Supplementary material for "Control of mitotic spindle disassembly through SUMO-targeted ubiquitylation of yeast kinesin-5": Table S3 Primers

| | Primer name | Sequence |
|---|---|---|
| 1190 | Cin8.K267R.for | 5'-AATCACTGCCGAATACCATCAgGCAACAGTATCAACAACAACA-3' |
| 1191 | Cin8.K267R.rev | 5'-TTAGTGACGGCTTATGGTAGTcCGTTGTCATAGTTGTTGTTGT-3' |
| 1220 | Cin8.K504R.for | 5'-CTATGGAATTAGCAAAGATTAgATCCGATTTACTCTCTACAAA-3' |
| 1221 | Cin8.K504R.rev | 5'-TTTGTAGAGAGTAAATCGGATCTAATCTTTGCTAATTCCATAG-3 |
| 1222 | Cin8.K669R.for | 5'-AACCAAATCTTGATATGATCAgAAATGAAGTACTGACTCTTAT-3' |
| 1223 | Cin8.K669R.rev | 5'-ATAAGAGTCAGTACTTCATTTCTGATCATATCAAGATTTGGTT-3' |
| 1381 | Cin8.K854R.for | 5'-GAACCCAAGCGCGTTAGATGGGAAAATTCATTTG-3' |
| 1382 | Cin8.K854R.rev | 5'-CAAATGAATTTTCCCATCTAACGCGCTTGGGTTC-3' |
| 1393 | Cin8.1036.F | 5'-GAGAAAAATGTTAAGGATTGAACCCGG-3' |
| 1394 | Cin8.1036.R | 5'-CCGGGTTCAATCCTTAACATTTTTCTC-3' |
| 1387 | Cin8.KED.for | 5'-CTATGGTAACAAGGAAGACGCTACCAAAGACG-3' |
| 1388 | Cin8.KED.rev | 5'-CGTCTTTGGTAGCGTCTTCCTTGTTACCATAG-3' |
